## Supplementary Methods for "Single molecule imaging and modelling of mRNA decay dynamics in the *Drosophila* embryo"

### Supplementary note: filtering of the genes, model formulation and Bayesian inference with MCMC

September 28, 2022

This supplementary file provides detailed information on the filtering scheme, the transcriptional regulation model with Gaussian processes used in this paper, and inference methods, including Markov Chain Monte Carlo sampling.

### Filtering of differentially expressed genes

We used the approach introduced in Kalaitzis and Lawrence (2011) to filter out the genes which do not exhibit time-dependent dynamics. We used two Gaussian processes regressions of the form

$$y = f(t) + \epsilon, \tag{1}$$

where $f(\cdot)$ is a Gaussian process and $\epsilon$ is Gaussian noise with zero mean and variance $\sigma^2$. We fit two models to the data: 1) a Gaussian processes regression with GP defined by a noise kernel, i.e. $f(x) \sim GP(0, K_{noise})$; 2) a Gaussian process regression with GP defined by an RBF kernel, i.e. $f(x) \sim GP(0, K_{RBF})$. We kept only the genes which exhibited the dynamics when comparing the null model with no time-dependence (noise kernel) and the model with time-dependence (RBF kernel).

### Model formulation through differential equations

Gene transcription is a complex process that depends on multiple factors. The dynamics of mRNA in time, $m(t)$, is often assumed to be determined by the level of pre-mRNA (approximated by the pol-II time series), $p(t)$, splicing efficiency $S$ and degradation of mRNA itself defined in our model through the parameter $D$. Thus, the dynamics of mRNA largely depend on the accumulation of pre-mRNA and the relation between splicing efficiency and degradation of mRNA. In Lawrence et al. (2007) and Honkela et al. (2015) pre-mRNA is modeled non-parametrically as a Gaussian process (GP). Let $f(t) \sim GP(0, \Sigma)$, where $\Sigma$ is defined by RBF kernel. Then pre-mRNA is assumed to be a Gaussian process

$$p(t) = f(t). \tag{2}$$

The dynamics of mRNA follows an ordinary differential equation (ODE)

$$\frac{dm}{dt} = B + Sp(t) - Dm(t), \tag{3}$$

where $B$ is the basal transcription rate of a gene, and the initial baseline expression level is assumed to be $m(0) = B/D$. $S$ is the sensitivity of a gene to the transcription factor. The solution to the differential Equation (3) reads

$$m(t) = \frac{B}{D} + Se^{-D\times t} \int_0^t f(u)e^{D\times u} du. \tag{4}$$

Note, that in Equation (4) the dependence of $m(t)$ on GP is linear since integration is a linear operation. It follows that $m(\cdot)$ is also a Gaussian process, albeit with a different kernel. For more detailed derivations of this kernel we refer to Lawrence et al. (2007) and Honkela et al. (2015), while we discuss the exact kernel function and implementation of the GP regression following described ODEs in the next section.

### Observational model with noise and GPflow implementation

We assume an observational model with noise, thus in practice, we observe $y^m(t)$ and $y^p(t)$ defined as

$$y^m(t) = m(t) + \epsilon^m(t), \tag{5}$$

$$y^p(t) = p(t) + \epsilon^p(t), \tag{6}$$

where the noise terms $\epsilon^m(t)$ and $\epsilon^p(t)$ are assumed to be Gaussian with mean zero and different variances. We assume the following kernel for $x^p(t)$:

$$k_p(t, t') = V \exp\left(-\frac{(t-t')^2}{l^2}\right). \tag{7}$$

The kernel for $m(\cdot)$ and cross-covariance between $m(\cdot)$ and $p(\cdot)$ are derived in Lawrence et al. (2007). The kernel for $m(\cdot)$ reads

$$k_m(t, t') = VS^2 \frac{\sqrt{\pi} l}{2} \left[h(t', t) + h(t, t')\right] \tag{8}$$

where

$$\begin{aligned} h(t', t) = \frac{\exp(\gamma)^2}{2D} \Bigg\{ & \exp[-D(t'-t)] \left[ erf\left(\frac{t'-t}{l} - \gamma\right) + erf\left(\frac{t}{l} + \gamma\right)\right] \\ & - \exp[-(Dt' + D)] \left[ erf(\frac{t'}{l} - \gamma) + erf(\gamma)\right], \end{aligned} \tag{9}$$

where $\gamma = \frac{Dl}{2}$.
The cross-covariance between $m(\cdot)$ and $p(\cdot)$ is

$$k_{mp}(t', t) = V \frac{\sqrt{\pi} lS}{2} \exp(\gamma)^2 \exp\left[-D(t'-t)\right] \left[ erf(\frac{t'-t}{l} - \gamma) + erf(\frac{t}{l} + \gamma)\right]. \tag{10}$$

We write the complete GP regression model by combining the data for mRNA and pre-mRNA into a joint vector $\boldsymbol{y} = (\boldsymbol{y^m}, \boldsymbol{y^p}) \sim GP(0, K_y + K_{noise})$, where joint covariance matrices for ODE-like dynamics and noise terms are given by

$$K_y = \begin{pmatrix} K_m & K_{mp} \\ K_{pm} & K_p \end{pmatrix}, \quad K_{noise} = \begin{pmatrix} \sigma_m^2 I & 0 \\ 0 & \sigma_p^2 I \end{pmatrix}. \tag{11}$$

We use GPflow library Matthews et al. (2017) for implementation, the block kernel described in Equation (11) is coded manually, but the routines for the optimization of the parameters are standard.

### Marginal likelihood

The marginal likelihood of the model is Gaussian and is used to optimize model parameters. For numerical stability, log-marginal likelihood is used:

$$\log p(y|t,\theta) = -\frac{1}{2}y^T K_y^{-1} y - \frac{1}{2}\log|K| - \frac{N}{2}\log 2\pi, \tag{12}$$

Then Metropolis-adjusted Langevin algorithm, initialized in optimized parameter values, is used to sample from the posterior distribution.

### Markov Chain Monte Carlo sampling from the posterior distribution

We use the Metropolis-adjusted Langevin algorithm (MALA) sampling scheme for fully Bayesian inference of the genes of interest. Using MALA we obtain posterior samples to quantify uncertainty about the parameters of the model. MALA is a gradient-based MCMC algorithm that exploits Langevin dynamics for the proposal mechanism. Generally, this allows obtaining a higher acceptance rate while making large proposal steps and exploring parameter space sufficiently well.

We use the implementation of MALA from the Tensorflow probability library while the model itself is implemented in GPflow library. For the sampling all of the parameters are transformed to the unconstrained space using logistic transformation.

The following prior distributions were chosen for the model parameters: $S \sim Gamma(2, 2)$, $D \sim Gamma(2, 2)$, $\sigma_m^2 \sim Gamma(\alpha_m, \beta_m)$ and $\sigma_p^2 \sim Gamma(\alpha_p, \beta_p)$, where hyperparameters $\alpha_m$, $\beta_m$, $\alpha_p$, $\beta_p$ depend on the empirical mean ans standard deviation.

We run MALA with different number of step sizes $[10^{-5}, 10^{-4}, 10^{-3}, 0.003, 0.005, 0.01, 0.03, 0.05, 0.07, 0.1, 0.15, 0.2, 0.25, 0.3, 0.35, 0.5, 1]$ and pick the one which results in higher acceptance rate/-better mixing of the posterior distribution.

### Illustrative examples

In this section, we use two sets of examples to demonstrate the performance of the model. In the first set of examples, we simulated five data sets with different degradation rates – 0.003, 0.008, 0.01, 0.02, 0.05 – which correspond to 231.0, 86.6, 69.3, 34.6, 13.8 min half-lives respectively. Figures S1**a**, **c**, **e**, **g**, **i** illustrate simulated data and fitted models and Figures S1 **b**, **d**, **f**, **h**, **j** illustrate corresponding posterior estimates for the degradation parameter obtained with

MCMC and with maximum a posteriori estimation. For all of the settings, the true degradation parameter is inferred accurately using both the maximum a posteriori approach and MCMC sampling.

In the second set of examples, we focus only on small degradation rates – 0.008, 0.003, 0.001, 0.0005 – which correspond to 86.6, 231.0, 693.1, and 1386.2min half-lives, respectively. We investigate the limit of the temporal scale of the half-lives, which is identifiable by the model. Figures S2 **a**, **b** , **c**, **d** illustrate model fits for the small degradation examples and the estimates of the degradation obtained with maximum a posteriori estimation. In Figures S2**a** and **b** the degradation rates, 0.008 and 0.003, are estimated well and are still identifiable. However, in Figures S2**c** and **d** the model cannot distinguish between degradation parameters 0.001 and 0.0005 (693.1min and 1386.2min half-lives respectively). When the degradation is zero, the half-life is practically indistinguishable from infinity, and thus, for small degradations (large half-lives), the parameter becomes non-identifiable.

In both sets of examples, other model parameters for data generation were fixed to: $S = 0.3$, $V = 69.0$, $l = 33.0$, $\sigma_m^2 = 50.5$, $\sigma_p^2 = 3.5$.

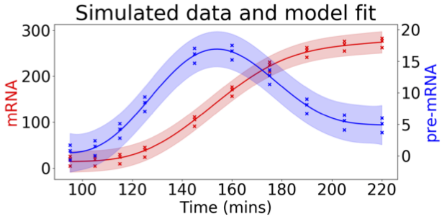


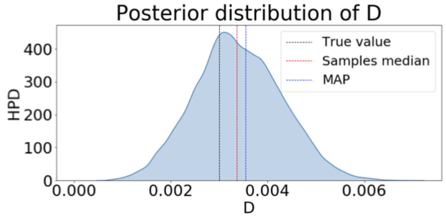


**a** $D$=0.003, half-life=231.0 min **b** $D_{map} = 0.0035$, $D_{med} = 0.0033$

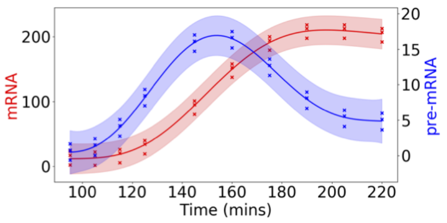


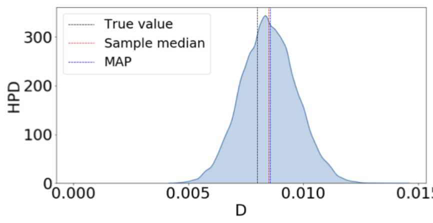


**c** $D$=0.008, half-life=86.6 min **d** $D_{map} = 0.0085$ , $D_{med} = 0.0084$

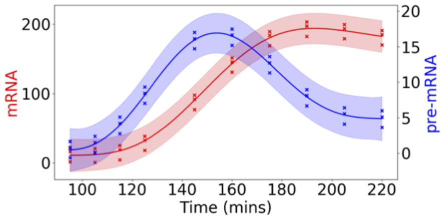


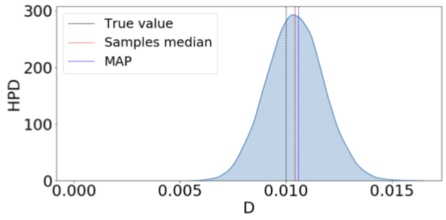


**e** $D$=0.01 half-life=69.3 min **f** $D_{map} = 0.01$ , $D_{med} = 0.01$

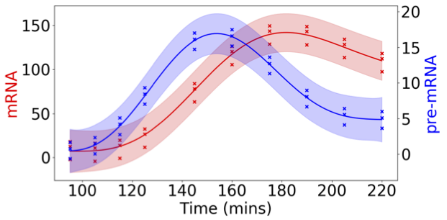


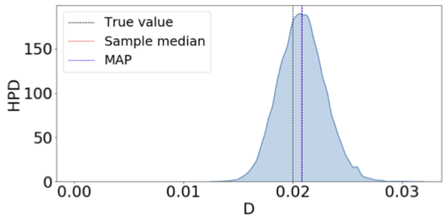


**g** $D$=0.02, half-life=34.6 min **h** $D_{map} = 0.02$ , $D_{med} = 0.02$

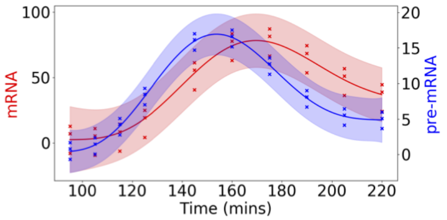


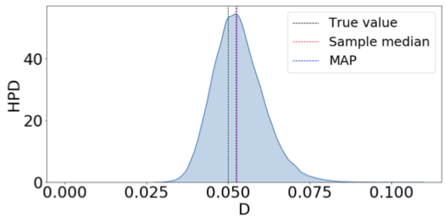


**i** $D$=0.05, half-life=13.8 min **j** $D_{map} = 0.052$ , $D_{med} = 0.052$



Figure S1: *Left column*: simulated data and model fits of transcriptional regulation model. *Right column*: posterior density estimates for the degradation parameters, where $D_{map}$ indicates maximum a posteriori estimate and $D_{med}$ indicates MCMC sample median.

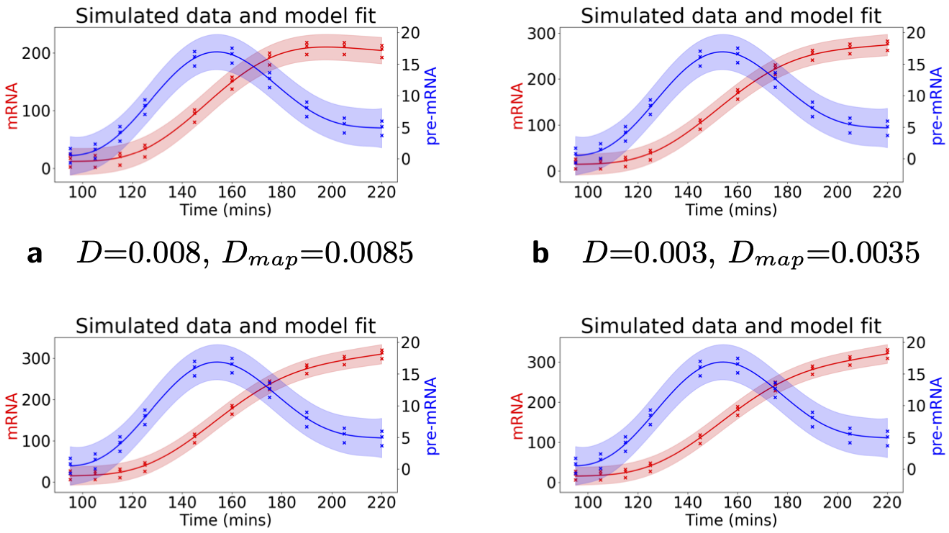


**a** $D$=0.008, $D_{map}$=0.0085

**b** $D$=0.003, $D_{map}$=0.0035

**c** $D$=0.001, $D_{map}$=0.0016

**d** $D$=0.0005, $D_{map}$=0.0012

Figure S2: Examples of model fits and maximum a posteriori estimates ($D_{map}$) for small $D$ values/long half-lives. **a** $D = 0.008$ corresponds to 86.6min half-life. **b** $D = 0.003$ corresponds to 231.0min half-life. **c** $D = 0.001$ corresponds to 693.1min half-life. **d** $D = 0.0005$ corresponds to 1386.2min half-life. When $D$ approaches zero, it becomes unidentifiable.
